# Iron can be microbially extracted from Lunar and Martian regolith simulants and 3D printed into tough structural materials

**DOI:** 10.1101/2020.11.15.382614

**Authors:** Sofie M. Castelein, Tom F. Aarts, Juergen Schleppi, Ruud Hendrikx, Amarante J. Böttger, Dominik Benz, Maude Marechal, Advenit Makaya, Stan J. Brouns, Martin Schwentenwein, Anne S. Meyer, Benjamin A.E. Lehner

## Abstract

*In-situ* resource utilization (ISRU) is increasingly acknowledged as an essential requirement for the construction of sustainable extra-terrestrial colonies. Even with decreasing launch costs, the ultimate goal of establishing colonies must be the usage of resources found at the destination of interest. Typical approaches towards ISRU are often constrained by the mass and energy requirements of transporting processing machineries, such as rovers and massive reactors, and the vast amount of consumables needed. Application of self-reproducing bacteria for the extraction of resources is a promising approach to avoid these pitfalls. In this work, the bacterium *Shewanella oneidensis* was used to reduce three different types of Lunar and Martian regolith simulants, allowing for the magnetic extraction of iron-rich materials. The quantity of bacterially extracted material was up to 5.8 times higher and the total iron concentration was up to 43.6% higher in comparison to untreated material. The materials were 3D printed into cylinders and the mechanical properties were tested, resulting in a 396 ± 115% improvement in compressive strength in the bacterially treated samples. This work demonstrates a proof of concept for the on-demand production of construction and replacement parts in space exploration.

## Introduction

For the first time since the Apollo missions to the Moon, space agencies and commercial partners from all over the world are dedicated to send humans beyond Low Earth Orbit (LEO). The most famous examples are the European Space Agency (ESA), who have announced their plan to build an international Moon village [1]; the National Aeronautics and Space Administration (NASA), who plan to send the first human to Mars [2]; and the American company SpaceX, who published their vision of realizing a human settlement on Mars [3].

A human outpost on another celestial body needs to fulfill several requirements. Transportation, shelter, supplies, waste removal, hazard protection, and power sources have to be provided to ensure its viability [4]. Every space exploration endeavor to date, however, has used resources originating from Earth, which leads to high transportation costs and, ultimately, a high degree of dependence on Earth resources. The two most common solutions to tackle the cost issue are to reduce the payload weight and to minimize the launching costs [5]. However, in the long run sustainability, earth independence, and economic feasibility can only be ensured via the direct usage of resources found in space, also called *in-situ* resource utilization (ISRU) [6].

Traditionally, research in the sector of ISRU has focused on investigating technical processes already in use on Earth and how they may be implemented in the harsh environment of space [7]. Such technological approaches that have survived in the free market over an extended period have proven themselves to be successful in this highly selective process. Current Earth technologies have been optimized and tested extensively and may also be suitable for adaptation to the environment on other celestial bodies [7]. The same microwaves which are used for heating food could, for instance, become a key technology to sinter building blocks for a possible extraterrestrial outpost [8].

When considering *in*-*situ* mining of building materials, direct adaptation of current mechanochemical approaches to space applications has a major disadvantage: the need for outsized factories and machinery. One novel solution to this problem is the usage of microbes to allow ISRU. These organisms can be produced in large quantities, only requiring water and an easily transportable growth medium [5]. Microbes can be successfully used in many different techniques to produce [9], extract [10], and pattern [11] material in a manner that is similar to the equivalent mechanochemical processes. Microbial production processes do not require sophisticated factories, high energy investment, or toxic chemicals [4]. Therefore, bacterial methodologies are increasingly being applied to terrestrial applications [12–15] and may be even more valuable for space exploration and colonialization, where resupply and a limiting initial amount of materials are major constraints [4, 16]. The research presented here aims to develop a new approach to extract iron from Martian regolith and use it in 3D printing applications (Fig. 1).

**Fig. 1.**
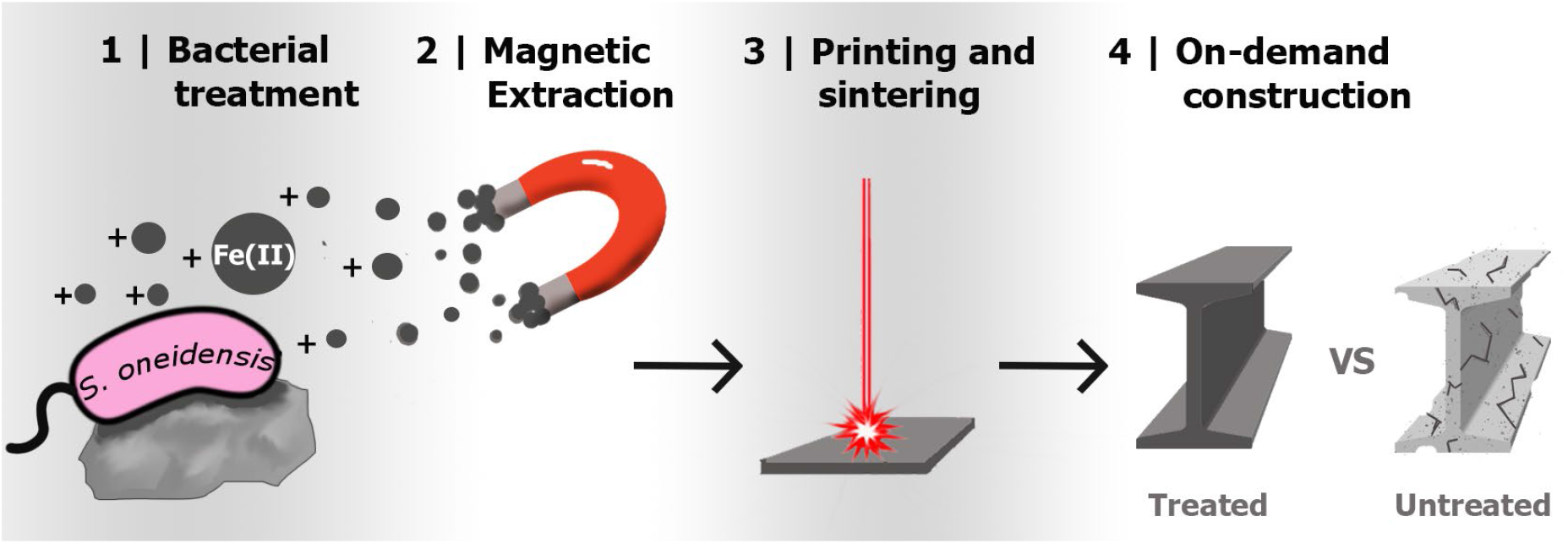
Conceptual flow of the microbial biomining process. Bacterial treatment (1) makes the material more magnetic by reducing iron. A magnet is applied to extract the magnetic material (2), and those particles are 3D printed and sintered (3). The material is then ready for construction, maintenance, and repair applications (4).

The bacterium *Shewanella oneidensis,* which was shown to be feasible for space-based biomining applications [17], can reduce a variety of metal oxides under both aerobic and anaerobic conditions [18, 19]. Martian regolith contains a high concentration of iron in the form of Fe(III) [20] that can be reduced to magnetite by *S. oneidensis* [21], which can subsequently be extracted magnetically. This work has developed a new methodology for the successful extraction and utilization of iron from several different regolith types by applying *S. oneidensis*, magnetism, and additive manufacturing (Fig. 1). Our results indicate that none of three tested regolith simulants are toxic to *Shewanella* under either aerobic or anaerobic conditions. Anaerobic iron extraction of JSC-Mars1 regolith simulant with minimal growth medium resulted in a 5.8-fold increase in the quantity of magnetically extractable iron and a total iron concentration that was 13.6% higher than in the starting material. The crude extracted iron was suitable for 3D printing using lithography-based ceramic manufacturing. Together, our findings provide the first proof-of-principle of bio-mediated *in-situ* iron extraction from Martian regolith simulant of a quality sufficient for on-demand production of infrastructure materials as well as replacement parts for a future space colony.

## Results & Discussion

### Compositions and limitations of regolith simulants

The elemental and mineral compositions of Martian regolith simulants, in comparison to actual Martian regolith, are of critical importance to make conclusions regarding the applicability of our regolith extraction methodology on Mars. X-ray fluorescence (XRF) measurements were performed on two Lunar Mare and one Martian regolith simulant to analyze their elemental compositions. Comparison of the two Lunar Mare simulants EAC-1 and JSC-2A with the average Martian regolith values (Tab. 1) [22] revealed that both contained similar oxide concentrations that resembled that of the average Martian soil. In contrast, the Martian regolith simulant JSC-Mars1 exhibited several differences in its elemental compositions compared to the average elemental composition of the actual Martian soil, especially for SiO_2_, Al_2_O_3_, and MgO. However, since the actual composition on Martian soil varies between different locations on the surfaces of the celestial objects, it is helpful to use a number of different simulants with varying compositions to test for multiple different scenarios. In contrast to real Lunar regolith, the simulants also contain Fe(III) and share key properties with Martian regolith, making them excellent candidates for our experiments.

**Tab. 1.** Average major metal oxide composition (wt%) at different Mars locations [26, 27, 31] and the composition of three different regolith simulant types obtained via X-ray fluorescence (XRF) (JSC-Mars1 [28], JSC-2 [29], EAC-1 [30]) as well as reported by the suppliers.

| Metal oxides (wt%) | Mars Viking 1 [26] | Mars Viking 2 [26] | Mars Pathfinder [27] | JSC-Mars 1 (Supplier) [28] | JSC-Mars1 (XRF) | JSC-2A (Supplier) [29] | JSC-2A (XRF) | EAC-1 (Supplier) [30] | EAC-1 (XRF) |
| --- | --- | --- | --- | --- | --- | --- | --- | --- | --- |
| SiO<sub>2</sub> | 43 | 43 | 44 | 43.5 | 42.19 | 47.5 | 44.89 | 43.7 | 43.58 |
| Al<sub>2</sub>O<sub>3</sub> | 7.3 | 7 | 7.5 | 23.3 | 23.48 | 15 | 19.61 | 12.6 | 11.45 |
| Fe<sub>2</sub>O<sub>3</sub> | 18.5 | 17.8 | 16.5 | 15.6 | 17.55 | 10.75 | 12.78 | 12 | 12.66 |
| MgO | 6 | 6 | 7 | 3.4 | 3.42 | 9 | 4.72 | 11.9 | 14.08 |
| CaO | 5.9 | 5.7 | 5.6 | 6.2 | 6.01 | 11 | 10.1 | 10.8 | 10.18 |
| K<sub>2</sub>O | <0.15 | <0.15 | 0.3 | 0.6 | 0.49 | 0.8 | 0.67 | 1.3 | 1.18 |
| TiO<sub>2</sub> | 0.66 | 0.56 | 1.1 | 3.8 | 3.80 | 2 | 2.23 | 2.4 | 2.15 |

The limitations of the different simulants used in our experiments will have an impact on the interpretation of our results. Since simulants are typically sourced and produced in varying ways, and the quality-assurance processes of manufacturing companies and agencies frequently differ from each other, the actual simulant quality will vary. Both Moon regolith simulants JSC-1A and EAC-1 have been mined from the surface of a volcanic basaltic deposit. However, JSC-1A was mined from a volcanic ash deposit in the San Francisco volcano field in Arizona (35°20’ N, 111°17’ W), which erupted only approximately 0.15 ± 0.03 million years ago [23], whereas EAC-1 was mined from a deposit in the Eifel region (50°41’N, 7°19’E), which is approximately 20 million years old. Although both simulants could have been in contact with water, air, and vegetation prior to being mined, the older age of EAC-1 increases the chance of exposure to processes that could change its composition. For example, EAC-1 shows alterations such as the presence of chlorite, which is usually found in igneous rocks and results from water interacting with pyroxene minerals. JSC-2A in contrast is produced from synthetic minerals [24] as the successor of JSC-1A and shares, therefore, the composition of JSC-1A without its environmental alterations due to air, water and vegetation. Therefore, we decided to use JSC-2A regolith rather than JSC-1A in our experiments.

The Martian regolith simulant JSC-Mars1 was sourced on the island of Hawaii from sieved palagonatized tephra ash from the 1843 Mauna Loa lava flow on Pu’u Nene (19°41’N, 155°29’W) [25]. Although all three regolith types were excavated close to the surface of the Earth, JSC-Mars1, at 175 years of age, is very young compared to the Lunar simulants EAC-1 and JSC-1A. Hence, few to no environmental alterations are expected in this simulant. Notably, actual Martian regolith contains sulfur trioxide (avg. 6.16%) and chlorine (avg. 0.68%), neither of which have been detected by the supplier or for the work presented here. The iron content of the Viking and Pathfinder Mars samples are best approximated by JSC-Mars1, while the aluminum content is closer to that of EAC-1 and the magnesium content is the closest to JSC-2A (Tab.1). These resemblances to real Martian regolith make all three regolith types (JSC-Mars1, EAC-1, and JSC-2A) good targets to study the interaction with *S. oneidensis*.

Equally as important as the elemental composition of regolith samples is the phase composition, which determines their accessibility for biological treatment with *S. oneidensis*. The mineral composition of the different regolith simulants was inspected by X-ray diffraction (XRD) analysis. The most abundant Martian minerals plagioclase, olivine, and pyroxene were detected in all three regolith simulants. EAC-1 showed the presence of nepheline and manganese iron oxide, both of which were not yet detected on Mars, and JSC-2A contained a small fraction of magnetite, similar to the results from Mars Curiosity (Tab. 2).

**Tab. 2.** X-ray diffraction analysis of the Mars Rocknest soil [20] and the three different regolith simulants. Minerals that were detected are denoted with “+”; minerals that were not detected are denoted with “−”. ^*^ These elements were only detected in the magnetically extracted fraction.

| Mineral | IUPAC nomenclature | Mars Curiosity Rocknest soil [20] | JSC-Mars1 (XRD) | JSC-2A (XRD) | EAC-1 (XRD) |
| --- | --- | --- | --- | --- | --- |
| **Plagioclase (Anorthite)** | (Ca<sub>0.57(13)</sub>Na<sub>0.43</sub>)(Al<sub>1.57</sub>Si<sub>2.43</sub>)O<sub>8</sub> | 40.8% | + | + | + |
| **Olivine (Forsterite)** | (Mg<sub>0.62(3)</sub>Fe<sub>0.38</sub>)<sub>2</sub>SiO<sub>4</sub> | 22.4% | + | + | + |
| **Pyroxene (Augite)** | (Mg<sub>0.88(10)</sub>Fe<sub>0.37</sub> Ca<sub>0.75(4)</sub>)Si<sub>2</sub>O<sub>6</sub> | 14.6% | + | (+)* | + |
| **Pyroxene (Pigeonite)** | (Mg<sub>1.13</sub>Fe<sub>0.68</sub>Ca<sub>0.19</sub>)Si<sub>2</sub>O<sub>6</sub> | 13.8% | - | - | - |
| **Magnetite** | Fe<sub>3</sub>O<sub>4</sub> | 2.1% | - | (+)* | - |
| **Anhydrite** | CaSO<sub>4</sub> | 1.5% | - | - | - |
| **Quartz** | SiO<sub>4</sub> | 1.4% | - | - | - |
| **Manganese Iron Oxide** | Mn<sub>0.43</sub>Fe<sub>2.57</sub>O<sub>4</sub> | - | - | - | + |
| **Nepheline** | $(K_{0.69}Na_{3.03}Ca_{0.03}Fe_{0.04}Al_{3.75}Si_{0.21})(SiO_4)_4$ | - | - | - | + |

Comparison of all three potential regolith types with the Martian regolith measurements revealed strong similarities in both mineral and elemental composition between the regolith simulants and the Martian target regolith. The Lunar regolith simulants are therefore suitable to use for our bio-based method.

### Toxicity of Regolith Simulants in aerobic and anaerobic conditions

To acquire a working concentration of regolith for our bacterial treatment, the toxicity of the Lunar and Martian regolith simulants for *S. oneidensis* was determined at a range of regolith concentrations under the presence (aerobic) and absence (anaerobic) of oxygen. *S. oneidensis* was grown for 48 hours in a rich tryptic soy broth (TSB) under aerobic (Fig. 2A, C, E) or anaerobic (Fig. 2B, D, F) conditions. Colony forming units (CFU) and optical density (O.D._600_) were measured to examine the effect of increasing regolith simulant concentrations (0 g/L, 0.1 g/L, 1 g/L, 10 g/L and 100 g/L) on both bacterial viability and growth. We tested the Martian regolith simulant JSC-Mars1 (Fig. 2A, B) and two Lunar regolith simulants: EAC-1 (Fig. 2C,D), JSC-2A (Fig. 2E,F). The CFUs of the samples were evaluated at three different time points (0, 24, and 48 hours). All control samples, which contained regolith but no bacterial cells, contained 0 CFU/μL at all time points. The O.D._600_ was measured every 5 minutes, and the optical density of control samples containing regolith but no *S. oneidensis* was subtracted from each data point.

**Fig. 2.**
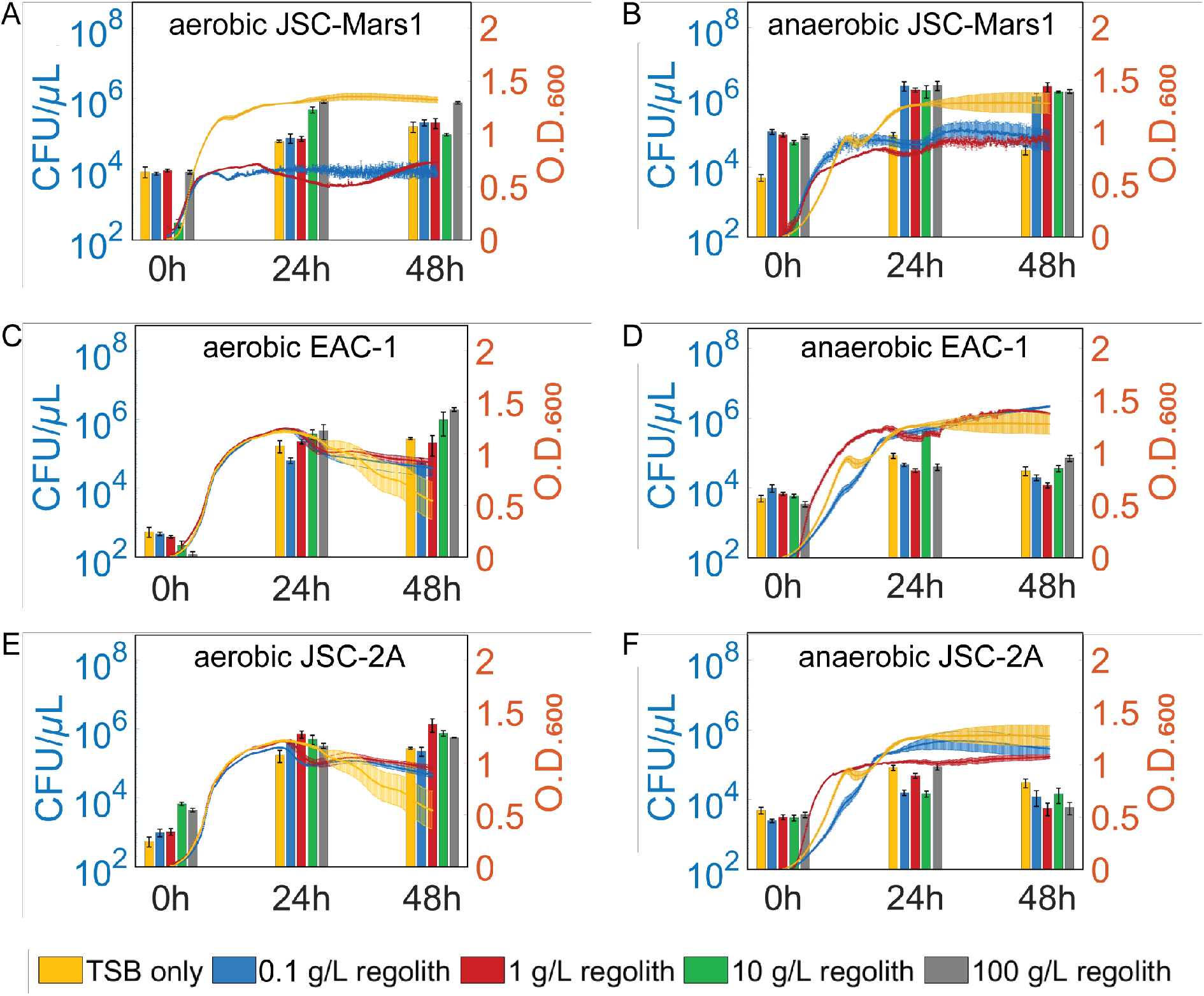
Evaluation of the toxicity of different Lunar and Martian regolith simulants to *S. oneidensis* under aerobic and anaerobic conditions. Colony forming units (CFU/μL) are depicted on the left axis and the O.D._600_ measurements on the right axis. The yellow, blue, red, green, and grey bars represent regolith concentrations of 0, 0.1, 1, 10, and 100 g/L respectively. Optical density at 600 nm of control samples without *S. oneidensis* were subtracted from the O.D._600_ measurements. The error bars represent the standard error of the mean. Aerobic growth behavior of *Shewanella oneidensis* was measured in the presence of JSC-Mars1 (A), EAC-1 (C) and JSC-2A (E), and anaerobic growth of *Shewanella oneidensis* was measured in the presence of JSC-Mars1 (B), EAC-1 (D) and JSC-2A (F).

*S. oneidensis* showed little or no reduction in maximal CFU concentration or O.D._600_ absorbance upon exposure to increasing concentrations of the different regolith types. The exception to this trend was a decrease in O.D._600_ absorbance that was observed upon exposure to increasing regolith concentrations in bacteria exposed to JSC-Mars1 compared to the no regolith control. This result may reflect the difficulty in obtaining reliable absorbance measurements in the presence of high iron concentrations due to the optical properties of iron. The bacteria exposed to JSC-2A and EAC-1 under aerobic conditions demonstrated higher maximal CFU concentrations compared to anaerobic conditions, indicating that the aerobic conditions resulted in a better growth environment. In contrast, bacteria exposed to JSC-Mars1 had a higher maximal CFU upon anaerobic growth, possibly due to the higher amount of Fe(III) that is available as a terminal electron acceptor in this regolith type. The starting CFUs of all anaerobic samples was slightly higher due to time required to transfer the samples from the anaerobic glove box to agar plates. Overall, the O.D._600_ and CFU results indicate similar growth behavior for bacteria exposed to different concentrations and types of regolith under the same environmental conditions. These results suggest low toxicity or affinity for any particular type or concentration of regolith towards *S. oneidensis*, with the exception of an affinity for JSC-Mars1 under anaerobic growth conditions.

### Magnetic extraction and Fe(II) content upon aerobic bacterial treatment

Construction material will be of utmost importance for any space colony, and magnetism would be a simple way to separate different ores. *S. oneidensis* can reduce Fe(III) to Fe(II) at its cell membrane together with a local pH increase, allowing for the precipitation of magnetite, a ferrimagnetic iron oxide. We hypothesized that this methodology might be applied to increase the quality and quantity of extractable magnetic material from regolith simulants. In order to produce more magnetic minerals within regolith and to concentrate the iron content, *S. oneidensis* was incubated over 168 hours with TSB medium and 10 g/L of EAC-1, JSC-2A, or JSC-Mars1 regolith simulant.

Each bacterially treated or untreated regolith simulant sample was assayed to determine both the change in amount of magnetic material and in aqueous Fe(II) concentration during treatment. To assess the amount of magnetic material at different timepoints, both with and without bacterial treatment, an aerobic magnetic extraction experiment was performed utilizing handhold neodymium magnets (Fig. 3A). A volume of 1 mL of the sample was pipetted into a cuvette. Then, a magnet was used to extract the magnetic fraction. After performing a washing step, the extracted material was transferred to a previously weighed cuvette. The weight difference of the material after drying was measured.

**Fig. 3.**
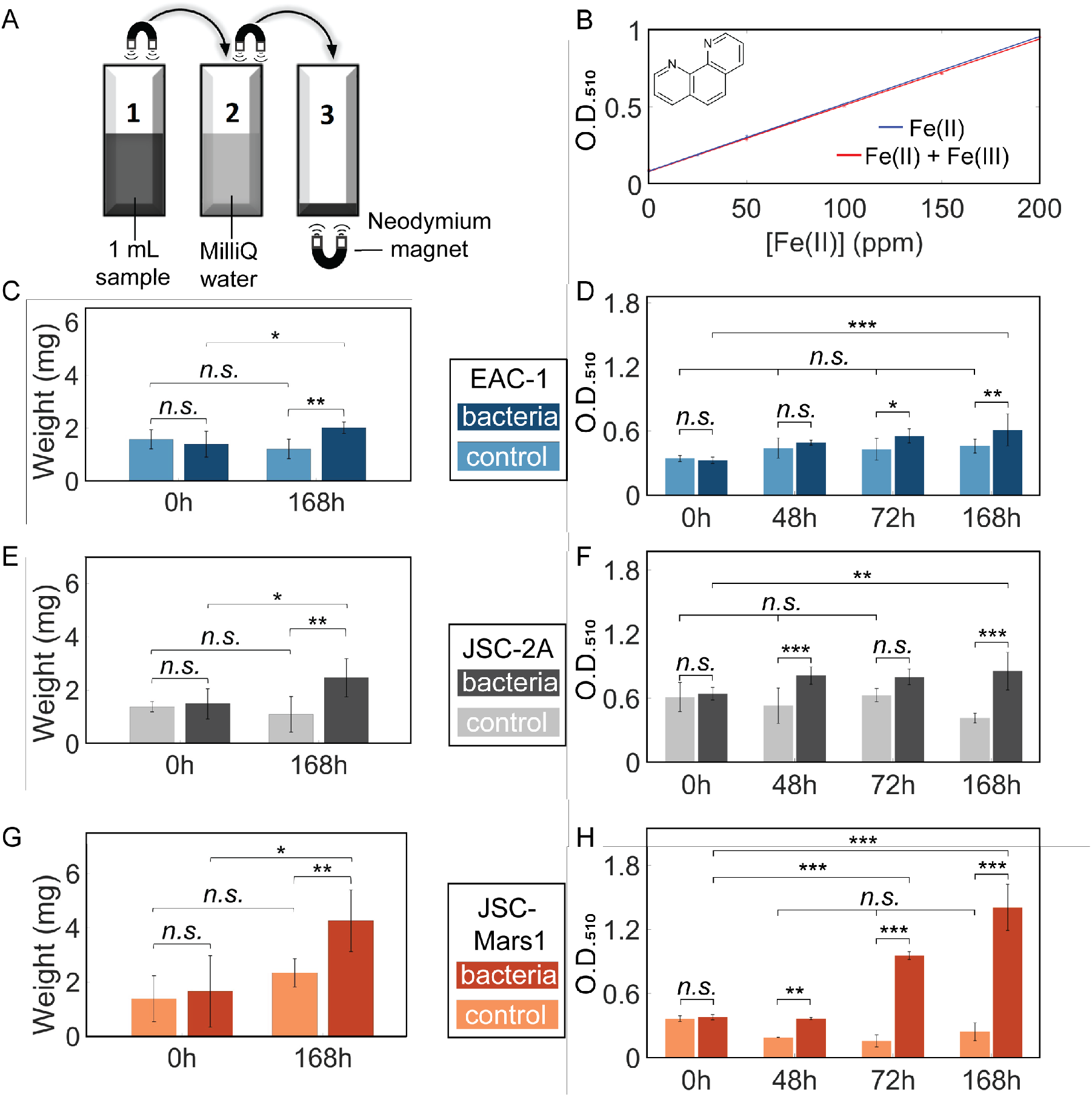
Aerobic extraction of magnetic material from EAC-1 (blue), JSC-2A (grey), and JSC-Mars1 (orange) regolith simulants. The darker color in each plot represents the bacterial sample, the lighter one the non-bacterial control. (A) The set-up for extracting magnetic material. In short, 1 mL sample (1) was pipetted into a cuvette, then the magnetic fraction was extracted via a magnet, washed in a MilliQ water (2), and finally dried in a previously weighed cuvette (3). The weight difference of the cuvette before and after this treatment represents the extracted magnetic material. (B) Absorbance of Fe(II) iron standards bound to 1,10-Phenanthroline in the absence and presence of Fe(III). O.D._510_ was measured for a range of 0 - 200 ppm Fe(II)_(aq)_ with or without a constant concentration of 100 ppm Fe(III)_(aq)_. (C, E, G) The weight of magnetically extracted material from samples containing 10g/L EAC-1 (C), JSC-2A (E), or JSC-Mars1 (G) solution with and without *S. oneidensis* after 0h and 168h. An increase of the magnetically extractable material was measured for all of the bacterially treated samples (n=6). (D, F, H) Absorbance of the colorimetric iron determination of 10 g/L EAC-1 (D), JSC-2A (F), and JSC-Mars1 (H) treated with aerobic *S. oneidensis* over 168 hours. A consistent increase of the Fe(II)_(aq)_ concentration was measured for all of the bacterially treated samples (n=9). Mean plus standard deviation is shown for the bar plots (One-way ANOVA with Tukey PostHoc test: n.s. > 0.05; * ≤ 0.05; ** ≤ 0.01; *** ≤ 0.001).

To determine the aqueous iron(II) concentration (Fe(II)_(aq)_) with and without bacterial treatment at different timepoints, a colorimetric assay was performed using 1,10-Phenanthroline as a complexing reagent for Fe(II) (Fig. 3B) [32]. An increase in Fe(II) bound to the complexing reagent can be measured via absorbance at O.D._510_. NaF was also added to complex Fe(III), thereby preventing interference with the absorbance measurement. This assay was performed on samples with a known iron concentration to generate a standard curve for Fe(II)_(aq)_ and also to test for any potential interference by Fe(III)_(aq)_ (Fig 3B). A Fe(II) standard curve was prepared by assaying ammonium iron(II) sulfate hexahydrate dissolved in TSB to a final concentration of 0, 50, 100, 150, or 200 ppm. A second standard curve was prepared by assaying samples containing the same Fe(II) concentrations as well as 100 ppm iron(III) citrate. A linear fit showed a good fit to a linear regression model (R^2^=0.9995). No significant difference was observed between the two curves (y_Fe(II)+Fe(III)_ = 4.3e-3*x + 8.0e-2; y_Fe(II)_ = 4.4e-3*x + 8.4e-2, One-way ANOVA, p = 0.99), indicating that no interference in the Fe(II) absorbance measurements was caused by the presence of up to 100 ppm Fe(III).

The amount of magnetically extracted material from EAC-1 increased significantly (Fig. 3C) from 1.4 ± 0.44 mg in the 0h sample to 2.0 ± 0.19 mg (1.4-fold) after 168 hours of incubation with *Shewanella* (p_S0h-S168h_ = 0.044), while there was no significant difference between the no-bacteria control at 0h and at 168h (One-way ANOVA with Tukey PostHoc test, n = 6, p_C0h-S0h_ = 0.83, p_C0h-C168h_ = 0.35). The 168h-incubated bacterially treated sample also contained significantly higher amounts of magnetic material than the 168h no-bacteria control (p_S168h-C168h_ = 0.0069).

Upon determination of the Fe(II) concentration of bacterially treated samples of EAC-1, the no-bacteria controls did not display any significant differences between the timepoints (Fig. 3D) (One-way ANOVA with Tukey PostHoc test, n=9, p > 0.05). The absorbance of the samples containing bacteria increased significantly within the first 48 hours (p_S0h-S48h_ = 0.0015) from 0.33 ± 0.029 A.U. to 0.40 ± 0.023 A.U., but was not significantly different from the control at 48h (p_C48h-S48h_ = 0.87). The Fe(II) concentration of the bacterially treated samples continued to increase throughout the remainder of the experiment to a final O.D._510_ of 0.61 ± 0.15 A.U., a 1.9-fold increase (p_S0h-S168h_ = 1e-8), corresponding to an aqueous iron concentration (Fe(II)_(aq)_) of 132.5 ± 17.5 ppm. The absorbance in the bacterial sample after 168 hours was also significantly greater than the control at 168h (p_C168h-S168h_ = 0.004), indicating a successful and significant reduction of Fe(III) to Fe(II) in EAC-1 after 72 and 168 hours of bacterial treatment.

In comparison to EAC-1, the amount of magnetically extracted material from JSC-2A regolith (Fig. 3E) increased more strongly between the 0h (1.5 ± 0.21 mg) and 168h (2.5 ± 0.27 mg) bacterially treated sample (p _S0h-S168h_ = 0.038). The 168h bacterially treated sample also contained significantly higher amounts of magnetically extracted material than the 0h (p_C0h-S168h_ = 0.019), and the 168h no-bacteria control (p_S168h-C168h_ = 0.0027). No significant differences were detected between the no-bacteria controls at 0h and 168h (p_C0h-C168h_ = 0.83), nor between the bacterial sample and the control at 0h (n = 6, one-way ANOVA with Tukey PostHoc test, p_S0h-C0h_ = 0.98). *S. oneidensis* was, therefore, able to increase the amount of magnetically extracted material in JSC-2A significantly (1.7-fold).

The starting Fe(II)_(aq)_ concentration in the JSC-2A bacterial samples (Fig. 3F) at 0h was double (0.64 ± 0.059 A.U.) that of EAC-1. The Fe(II) concentration in the bacterially treated sample was significantly higher than in the no bacteria control at both 48h (p_C48h-S48h_ = 3e-5) and 168h (p_S168h-C168h_ = 1e-8), but not at 72h (pC72h-S72h = 0.087). A significant increase in Fe(II) was observed between the bacterial sample at 0h and 48h (p_S0h-S48h_ = 0.035) as well as 168h (p_S0h-S168h_ = 0.0051) to a maximum of 0.85 ± 0.18 A.U. (1.3-fold), corresponding to a Fe(II)_(aq)_ concentration of 192.5 ± 25 ppm. However, the control at 168h was also significantly different from the control at 0h (p_C0h-C168h_ = 0.0086). The rest of the controls did not have any significant differences between each other at any timepoint (n = 9, one-way ANOVA with Tukey PostHoc test, p > 0.05). These results show that a significant reduction of Fe(III) to Fe(II) by *S. oneidensis* can also happen in the bacterially treated JSC-2A regolith simulant, but the increase in Fe(II) was smaller since JSC-2A contains less Fe(III) than EAC-1.

The JSC-Mars1 sample (Fig. 3G) showed the highest increase in magnetically extractable material (2.5-fold) between the 0h (1.7 ± 1.2 mg) and the 168h (4.3 ± 1.04 mg; p_S0h-S168h_ = 5.5e-05) bacterial sample among all regolith simulants tested. The magnetically extracted weight of the 168h bacterial sample was also significantly higher than the 168h control (p_S168h-C168h_ = 0.025) and the 0h control (p_C0h-S168h_ = 1.02e-5). There were no significant differences between the controls (p_C0h-C168h_ = 0.32), nor between the control and bacterial sample at 0h (One-way ANOVA with Tukey PostHoc test, p_S0h-C0h_ = 0.89).

The ferrous iron (Fe(II)_(aq)_) concentration of JSC-Mars1 regolith (Fig. 3H) was significantly higher in the 0h no-bacteria control compared to the 48h and 72h controls (p < 0.05), but not compared to the 168h control (p_C0h-C168h_ = 0.071). However, significant changes in the Fe(II)_(aq)_ concentration occurred in the bacterial JSC-Mars1 sample, which increased from an initial value of 0.38 ± 0.025 A.U. at 0h to 0.95 ± 0.036 A.U. after 72h (p_S0h-S72h_ = 1e-8) and to 1.4 ± 0.22 A.U. after 168h equaling a 3.7-fold increase and a total Fe(II)_(aq)_ concentration of 330 ± 35 ppm (p_S0h-S168h_ = 1e-8). For all timepoints besides 0h, the bacterial sample was also significantly different from its respective controls (One-way ANOVA with Tukey PostHoc test, p < 0.05). JSC-Mars1 was therefore the best candidate for bacterial treatment and magnetic extraction, showing the highest increase in extractable material and dissolved Fe(II)_(aq)_ concentration. This regolith simulant was therefore explored further to test the effect of anaerobic incubation with *S. oneidensis* on Fe(II) content and extraction of magnetic material.

### Magnetic extraction and Fe(II)_(aq)_ content upon anaerobic bacterial treatment

Anaerobic growth experiments were performed to assess the reduction capability of *S. oneidensis* without the availability of oxygen as additional electron acceptor. The first set of anaerobic experiments used rich TSB as a growth medium. Quantification of the magnetically extracted materials (Fig. 4A) displayed a significant increase after both 72h (4.0 ± 0.74 mg) and 168h (3.6 ± 1.4 mg) compared to the no-bacteria control samples and the 0h bacterial sample (1.5 ± 0.31 mg) (p_S72h-C72h_ = 3.3e-5, p S0h-S72h = 0.00021, p_S168h-C168h_ = 0.0059, p _S0h-S168h_ = 0.00061). The amount of material extracted from the bacterial samples increased 2.7-fold between 0h and 72h and 2.4-fold between 0h and 168h. No significant increase in extracted material was observed in the no-bacteria sample over time, nor between the bacterially treated samples at 72h and 168h (One-way ANOVA with Tukey PostHoc test, p > 0.05). Therefore, the amount of magnetically extracted material for JSC-Mars1 was similar after 168h for both the anaerobic and aerobic conditions, but unlike the aerobic experiments, the maximal yield was already reached after 72 hours upon anaerobic treatment.

**Fig. 4.**
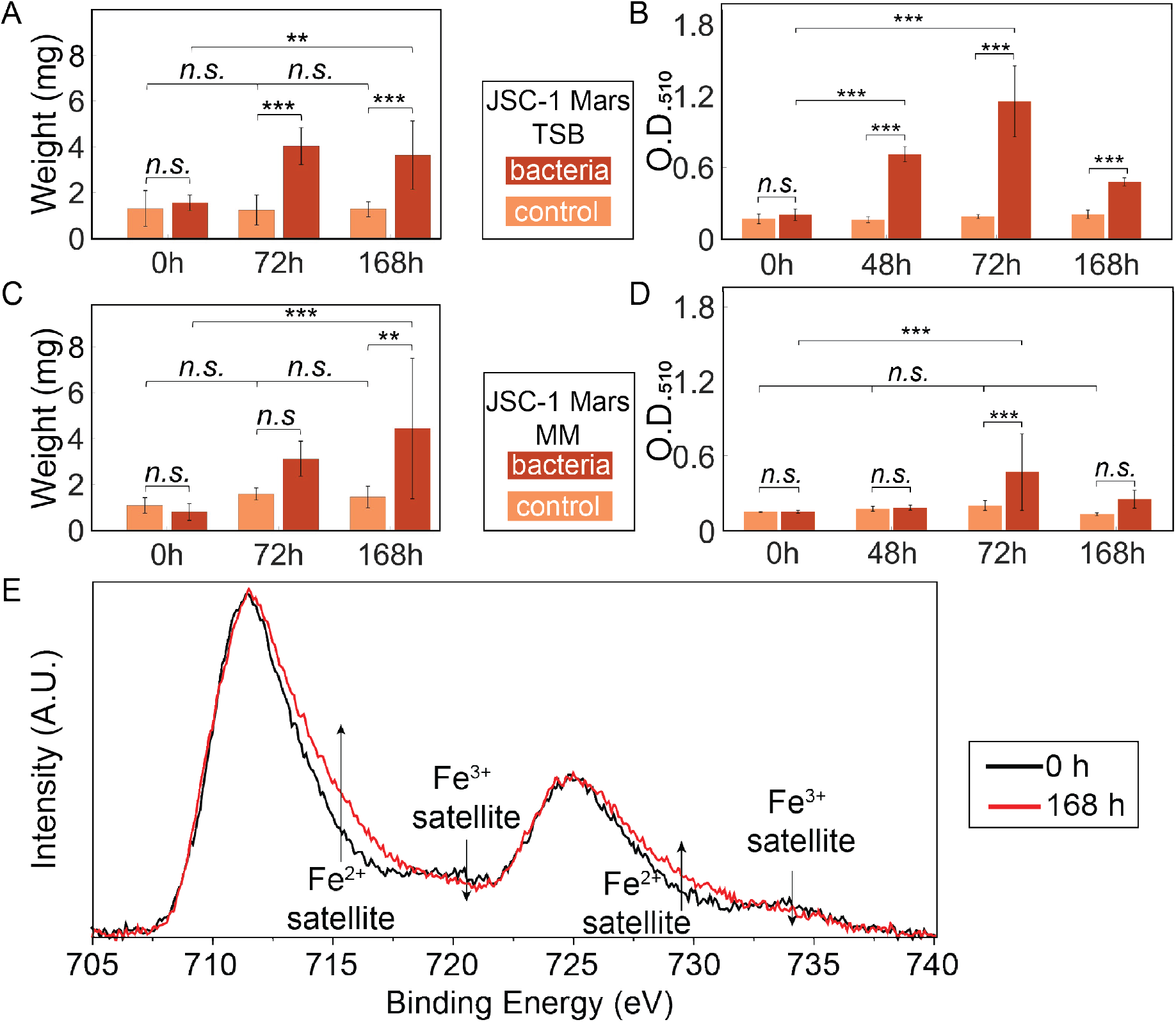
Anaerobic extraction of magnetic material from bacterially treated JSC-Mars1 regolith simulant. The amount of magnetically extracted material from a 10g/L JSC-Mars1 regolith simulant anaerobically incubated with (red) or without (orange) *Shewanella* bacteria in TSB medium (A) or defined minimal medium (MM) (C) after 0h, 72h, and 168h. The O.D._510_ values of the colorimetric assay to determine the Fe(II)_(aq)_ concentration of 10 g/L JSC-Mars1 regolith anaerobically incubated with (red) or without (orange) *Shewanella* bacteria in TSB medium (B) and defined minimal medium (D). The standard deviation is given for all bar plots (One-way ANOVA with Tukey PostHoc test: n.s. > 0.05; * ≤ 0.05; ** ≤ 0.01; *** ≤ 0.001). (E) X-ray photoelectron spectroscopy of iron in JSC-Mars1 regolith samples. A change is visible in the Fe(II) and Fe(III) satellite peaks between the 0h (black line) and the 168h (red line) timepoints of incubation with *S. oneidensis*.

The colorimetric assay to test the ferrous iron concentration (Fe(II)_(aq)_) displayed no significant differences between the absorbances of the control samples at any time point (Fig. 4B), nor between the bacterial sample and the control at 0h (One-way ANOVA with Tukey PostHoc test, p > 0.05). The bacterially treated sample showed a 3.6-fold, significant increase of Fe(II)_(aq)_ after 48h (0.71 ± 0.093 A.U.) and a 5.8-fold significant increase after 72h (1.16 ± 0.29 A.U.) compared to the 0h sample (0.20 ± 0.024 A.U.) (p_S48h-C48h_ = 1e-8, p_S0h-S48h_ = 1e-8, p_S72h-C72h_ = 1e-8, p _S0h-S72h_ = 1e-8). Interestingly, the Fe(II)_(aq)_ concentration decreased significantly after 168h (0.48 ± 0.032 A.U.) compared to the 72h bacterial sample (One-way ANOVA with Tukey PostHoc test, p_S168h-S72h_ = 1e-8). The greater yield and faster increase of Fe(II)_(aq)_ in the anaerobic JSC-Mars1 samples compared to the aerobic conditions indicates that the reduction of Fe(III) to Fe(II) was improved under anaerobic conditions.

JSC-Mars1 regolith simulant was also incubated anaerobically with *S. oneidensis* in a minimal, defined medium instead of the nutrient-rich TSB medium. A minimal medium holds the advantage that its nutrients can be fully utilized, which is essential to reduce the transportation weight for space applications. Quantification of the magnetically extracted material (Fig. 4C) showed no significant differences between any of the time points for the no-bacteria controls (p > 0.5). However, the bacterially treated, magnetically extracted material increased significantly after 72h to 3.1 ± 0.70 mg (3.8-fold) and 168h to 4.4 ± 2.8 mg (5.5-fold) in comparison to the 0h bacterially treated sample (0.8 ± 0.34 mg) and the no-bacteria control at 168h (p_S0h-S72h_ = 0.05, p_S168h-C168h_ = 0.0059, p _S0h-S168h_ = 0.00061). Unlike the anaerobic sample with TSB, no significant difference between the bacterially treated sample and the control after 72h was measured (One-way ANOVA with Tukey PostHoc test, p_S72h-C72h_ = 0.36). The absolute weight of the magnetically extracted material after 168h of anaerobic bacterial treatment of JSC-Mars1 in minimal defined medium (4.4 ± 2.8 mg) was comparable to the weight obtained from the anaerobic extractions in TSB (3.6 ± 1.4 mg) and the aerobic JSC-Mars1 treatment (4.3 ± 1.04 mg).

The Fe(II)_(aq)_ concentration in the colorimetric assay did not display significant differences between the no-bacteria controls at any timepoint (Fig. 4D), nor between the control and the bacterial sample at 0h (p > 0.05). A significant, 3.1-fold increase in Fe(II) to a maximum concentration of 97.5 ± 0.47 ppm was observed for the bacterial sample after 72h compared to 0h (p _S0h-S72h_ = 0.0082), followed by a significant decrease to 45 ± 1.75 ppm after 168h (p _S72h-S168h_ = 0.013). Only the 72h bacterially treated sample showed a significantly higher Fe(II)_(aq)_ concentration compared to its respective 72h no bacteria control (One-way ANOVA with Tukey PostHoc test, p _S72h-C72h_ = 0.00089). However, the increase in ferrous iron was lower than observed in the experiments using TSB. The experiments in a minimal defined medium showed a significant increase in both the Fe(II)_(aq)_ concentration as well as the magnetically extracted material after 72h and required the least resources, which makes this condition the preferred choice for space exploration.

We performed high-resolution X-ray photoelectron spectroscopy (XPS) measurements (Fig. 4E) to investigate the bacterial reduction of Fe(III) to Fe(II) in the anaerobically bacterially treated JSC-Mars1 samples after 0 and 168 hours. An increase in the XPS signal was observed in the 168-hour sample at approximately 715 eV and 728 eV, corresponding to Fe(II) satellite peaks. Additionally, the signals at 720 eV and 734 eV, corresponding to Fe(III) satellite peaks, showed a lower intensity for the 168h treated samples. Therefore, the XPS measurement confirmed the results of the colorimetric assay that anaerobic treatment of JSC-Mars1 regolith with *S. oneidensis* resulted in a successful reduction of Fe(III) to Fe(II).

### Iron and silicon concentration in the magnetically extracted materials

A higher iron oxide and lower silicon oxide concentration are critical to improve the electric as well as mechanical properties of bacterially treated regolith and decrease the melting point of the material [33]. We postulate that the magnetic treatment will concentrate iron and therefore improve the mechanical properties of the treated sample. To analyze the weight percent of iron oxides (Fig. 5A) and silicon oxides (Fig. 5B) in different regolith simulant samples, X-ray fluorescence measurements (XRF) were performed to evaluate the compositional changes of this treatment. *S. oneidensis* was incubated aerobically in the presence of regolith simulants for 168 hours, after which magnetic material was extracted by immersing a handheld neodymium magnet into the sample until no more magnetic material could be extracted. XRF measurements were then performed to determine the iron and silicon content of both the untreated regolith samples, to which no magnetic or bacterial methodology were applied, and the bacterially treated samples. The XRF data for our experimental samples can be directly compared to the values given by the supplier. The untreated regolith samples were found to have iron and silicon weight percentages equivalent to the values reported by the suppliers. In all three regolith types, an increase in the iron wt% (EAC-1 43.6%, JSC-2A 17.1%, JSC-Mars1 18.2%) and a decrease in the silicon wt% (EAC-1 −22.6%, JSC-2A −7.4%, JSC-Mars1 - 25.0%) were observed upon bacterial treatment and magnetic extraction compared to the untreated material. This alteration of the composition upon treatment indicates that not only the quantity but also the quality of extracted material for construction applications were improved through the application of *S. oneidensis.*

**Fig. 5.**
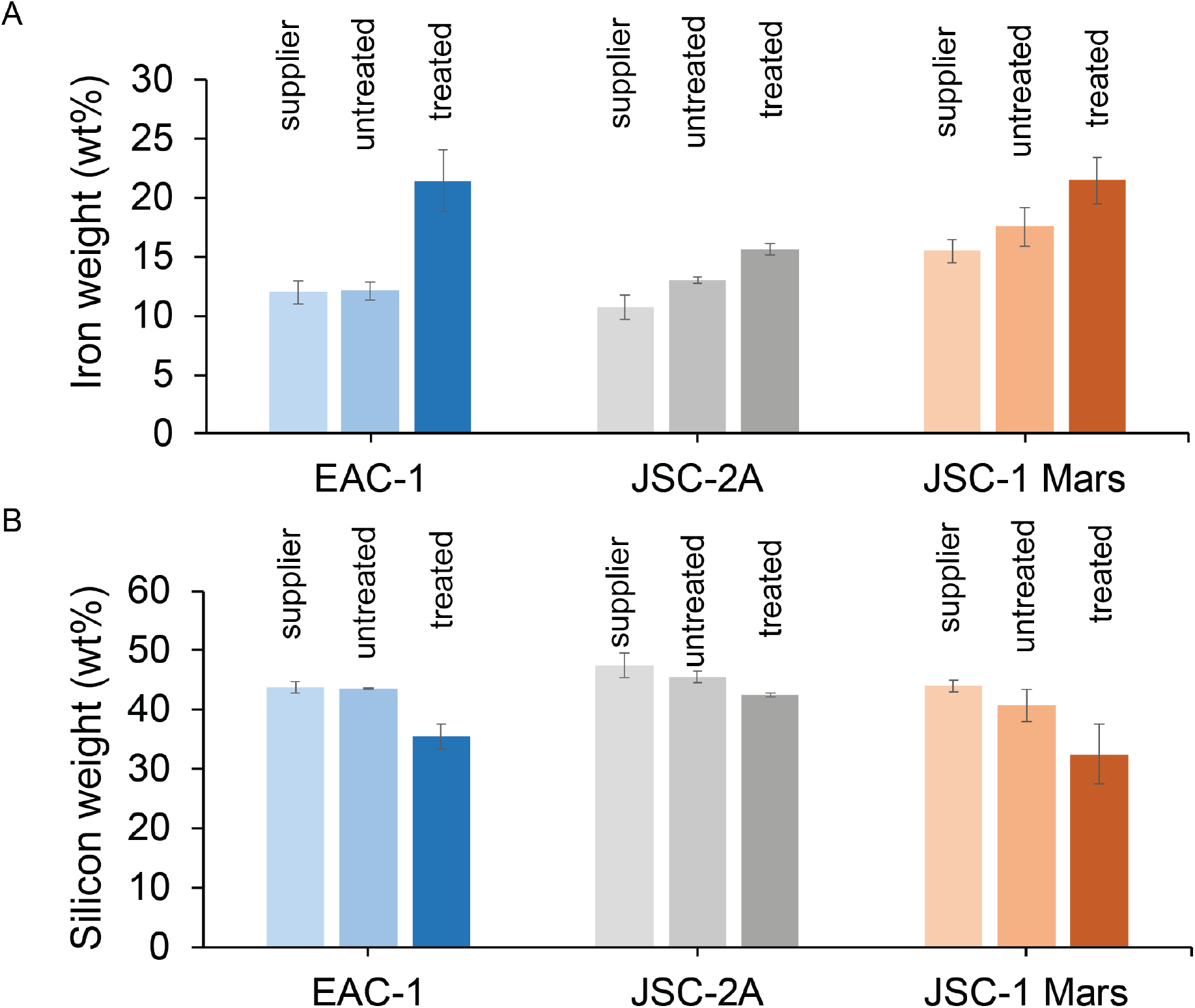
X-ray fluorescence spectroscopy analyzing the (A) iron concentration (wt%) and (B) silicon concentration (wt%) of the three different regolith simulants as given by the supplier (“supplier”, shown in bright colors); measured without treatment (untreated, in intermediate colors); or bacterially treated and magnetically extracted (treated, in dark colors). The different regolith types are indicated in blue for EAC-1, grey for JSC-2A, and red for JSC-Mars1. Error bars display the standard deviation.

### Ultimate Compressive Strength of 3D prints

To test whether bacterially and magnetically extracted regolith could produce a structural material with improved mechanical properties, 3D prints of treated and untreated JSC-2A were produced via lithography-based additive manufacturing followed by sintering (Fig. 6).

**Fig. 6.**
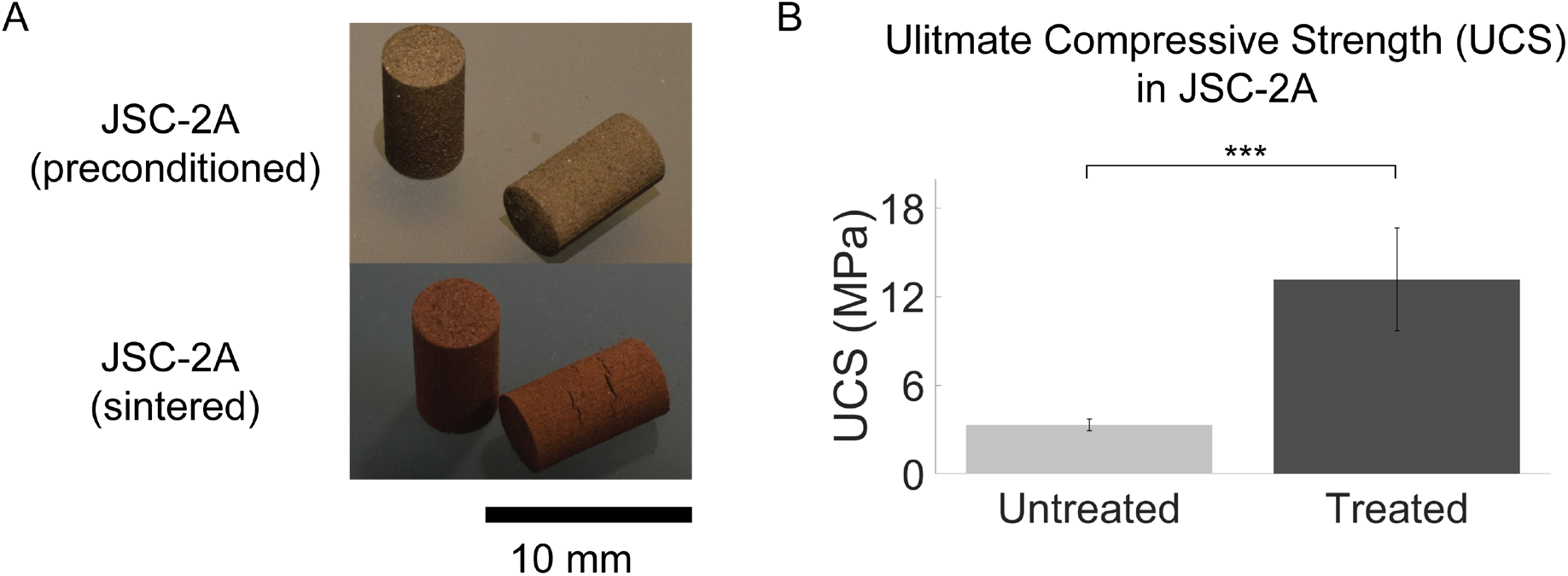
Lithography-based ceramic manufacturing (LCM) prints of JSC-2A regolith-based materials. (A) Lithography-based ceramic manufacturing of the regolith simulant JSC-2A (B) The ultimate compressive strength (UCS) of untreated (n=9) and the bacterially treated and magnetically extracted material (n=5). Error bars are displayed as standard deviation (Student’s t-test: n.s. > 0.05; * ≤ 0.05; ** ≤ 0.01; *** ≤ 0.001).

For the untreated samples, an average ultimate compressive strength of 3.33 MPa ± 0.39 MPa was measured (n=9). Strikingly, for the treated samples, an average ultimate compressive strength of 13.18 MPa ± 3.49 MPa was found (n=5), indicating a significant 400% increase of the ultimate compressive strength after bacterial treatment of the material (One-way ANOVA with Tukey PostHoc test, p = 1.7e-6). These results indicate that the sintered 3D prints produced from treated regolith are stronger than the ones produced from untreated regolith.

## Conclusion

In this work, we demonstrate an increase in the iron concentration and the quantity of magnetically extracted material upon bacterial treatment of three different regolith simulants. This increased iron concentration and quantity are both critical for applying our methodology to 3D printing applications, demonstrating an on-demand production work-flow for a space colony.

### Bacterial reduction of regolith simulants

*S. oneidensis* growth kinetics were not influenced by the presence of differing regolith simulant types or concentrations, which makes this bacterium an excellent candidate for ISRU operations. While its main purpose in this study is the reduction of Fe(III) to Fe(II) in a range of different regolith simulants, other activities of *S. oneidensis* might also be beneficial for iron extraction, including its reductive bioleaching activity of minerals [34] and its ability to generate locally increased pH values at the cell membrane, which is advantageous for the precipitation of magnetite [21].

The two Lunar regolith simulants JSC-2A and EAC-1 consist primarily of Fe(II)-containing minerals, making them a worse target for bacterial treatment with *S. oneidensis* than the Mars regolith simulant JSC-Mars1, which primarily contains Fe(III). Aerobic bacterial treatment of EAC-1 and JSC-2A samples showed a 1.9-fold and 1.3-fold increase in dissolved Fe(II)_(aq)_ respectively, compared to the higher 3.7-fold increase observed in bacterially treated JSC-Mars1. Remarkably, the quantity of extracted material, nevertheless, increased significantly by 1.7-fold in JSC-2A after 168 hours of bacterial treatment. The magnetic extraction of EAC-1 showed some significant increases in the bacterially treated sample, but contrasting trends were observed between the measured Fe(II)_(aq)_ concentrations (JSC-2A < EAC1) and the amount of magnetically extracted material (EAC-1 < JSC-2A) in bacterially treated samples. These results may indicate that not only the reduction, but also other bacterial functions (bioleaching, local pH) play an important role in the methodology.

The highest weight increase of magnetically extracted material (2.5-fold) and Fe(II)_(aq)_ concentration (3.7-fold) under aerobic conditions was measured in bacterially treated JSC-Mars1 samples. This result was likely due to differences in starting composition among the various regolith simulants, since JSC-Mars1 contains the highest Fe(III) concentration. Intriguingly, we were also able to successfully extract magnetic material from regolith simulants JSC-2A and EAC-1, which primarily contain Fe(II) rather than Fe(III), perhaps due to surface oxidation during their storage leading to increased levels of Fe(III). None of the regolith simulants was stored in an oxygen-free environment, making the outer surfaces of the materials particularly vulnerable to atmospheric oxidation.

The anaerobic bacterially induced reduction of Fe(III) in JSC-Mars1 with TSB as growth medium, compared to the aerobic experiments, resulted in an even further improved 5.8-fold increase of Fe(II)_(aq)_ after 72h. Interestingly, the Fe(II)_(aq)_ concentration of the bacterial sample was seen to decrease after 168h. The enhanced reduction and high concentration of reduced iron under anaerobic conditions could potentially indicate a faster precipitation of the produced magnetite, making the iron unavailable for the colorimetric assay.

Anaerobic bacterial reduction of JSC-Mars1 in the defined minimal medium produced a similar result, where the Fe(II) concentration reached a maximum increase of 3.1-fold at 72 hours. The overall lower values of iron reduction observed in minimal medium compared to TSB medium might be caused by lower concentration of Fe(III) ions in the defined medium compared to the rich TSB medium. A less successful reduction mechanism in the minimal medium under anaerobic conditions, due to starvation, is unlikely because of the 5.5-fold increase observed in magnetically extracted material from bacterially treated JSC-Mars1 regolith simulant after 168h, the highest increase in extracted material observed among all of our conditions. The anaerobic magnetic extraction in TSB medium was already at its maximum after 72h which indicates a faster reduction rate in rich medium under anaerobic conditions. The improved reduction and extraction in anaerobic conditions is likely due to the absence of oxygen as an alternative electron acceptor. In this environment, all electrons produced during the incubation with *S. oneidensis* are donated to the metal oxides of the regolith simulants as end terminal acceptor. Nevertheless, the maxima of extracted material of all three conditions in which JSC-Mars1 was bacterially treated (aerobically in TSB, anaerobically in TSB and minimal medium) do not significantly differ from each other. Therefore, reduction in the minimal medium under anaerobic conditions is the optimal candidate for space exploration, where the transport of additional nutrients or oxygen is a high burden on its feasibility.

The increase in iron and decrease in silicon concentration after the bacterial and magnetic extractions are essential quality factors for 3D printing applications. The ultimate compression strength was influenced by this factor, such that the bacterially and magnetically extracted 3D printed sample could withstand a pressure four times higher compared to the untreated one [35]. Additional 3D printing experiments coupled to material tests lead to even greater understanding of these properties.

Our hypothesis is that *S. oneidensis* primarily reduces the surface of the regolith simulant particles [36]. It is likely that bacterial reduction of Fe(III) to Fe(II) results in the appearance of more magnetic areas on the surface of regolith particles, so that an increased number of particles can be magnetically extracted. JSC-Mars1 has both a higher iron concentration as well as a higher Fe(III):Fe(II) ratio, presumably leading to an increased extraction yield after applying our methodology. Moreover, the JSC-Mars1 regolith simulant is expected to have a composition similar to many potential landing sites on Mars.

The methodology presented here shows the extraction of iron-rich material from different regolith simulants for 3D printing applications with improved mechanical properties compared to raw regolith. The building of sustainable habitats or replacement parts such as screws, airlocks and antennas will be all within the realm of possibility. This technology will enhance the *in-situ* resource utilization opportunities of humans using microorganisms and will, therefore, pave the way for future space exploration and Mars colonialization.

## Materials & methods

### Bacterial growth conditions

*Shewanella oneidensis* MR-1 (ATCC® 700550™) was inoculated into sterilized 32 g/L Tryptic Soy Broth (TSB) media and planktonically grown overnight under aerobic conditions at 30°C under continuous shaking (180-250 rpm) to an O.D._600_ of 0.5.

### Regolith pretreatment

All regolith simulant samples were sieved down to a maximum diameter of 63 μm. Only the particles smaller than 63 μm were used for further experiments.

### Preparation of anaerobic samples

Regolith was diluted to the desired concentration in TSB or defined minimal medium. The defined minimal medium was prepared by mixing 100 mM NaCl, 50 mM sodium 4-(2-hydroxyethyl)-1-piperazineethanesulphonic acid (HEPES), 7.5 mM NaOH, 16 mM NH_4_Cl, 1.3 mM KCl, 4.3 mM NaH_2_PO_4_·2H_2_O and 10 mL/L trace mineral supplement (ATCC® MD-TMS™) [37]. The defined minimal medium was autoclaved, after which it was supplemented with 10 mL/L vitamin solution (ATCC® MD-VS™) [38], 20 mg/L of L-arginine hydrochloride, 20 mg/L L-glutamate, 20 mg/L L-serine, and 20 mM lactate [39]. The samples were prepared in anaerobic culture tubes (Sigma Aldrich) and flushed 3x with 100% nitrogen gas to deplete the oxygen present in the solutions. After 24 hours, *Shewanella oneidensis* overnight cultures were added to the samples to a final O.D._600_ of 0.05. The samples were sealed with sterilized nontoxic chlorobutyl stoppers and aluminum lids (Sigma Aldrich).

### Toxicity of regolith simulants

Previously sterilized regolith simulant samples were sterilized again via autoclaving and diluted to the desired concentration in TSB. Overnight cultures of *Shewanella oneidensis* were added to the samples to a final O.D._600_ of 0.05. The bacterial growth in the samples was observed in a 96-well plate via optical density (O.D.) measurements in a plate reader (Tecan infinite M200 pro microplate reader) at a wavelength of 600 nm, during incubation at 30°C with continuous shaking (250 rpm) under aerobic or anaerobic conditions. Absorbance was measured every 5 minutes for 48 hours. The samples were measured in triplicate.

In parallel, the same samples were used to measure colony forming units (CFU). A logarithmic dilution curve was prepared of every sample to a 10^−7^ dilution and then plated out in triplicate onto LB agar plates at the 0, 6, 24, and 48-hour timepoints. Dilution was carried out using phosphate-buffered saline (PBS) solution. Samples were grown for 24 hours at room temperature (20-22°C), then individual colonies were counted.

### Fe(II)_(aq)_ concentration determination

Iron concentrations were determined based on a colorimetric method [40], which is a modification of the Phenanthroline method [32]. By dissolving 0.6 grams sodium fluoride (Sigma-Aldrich, 99%) in 28 mL MilliQ water and 0.57 mL sulfuric acid (99.999%), a complexing reagent was formed. An o-Phenanthroline solution was prepared mixing 30 μL of hydrochloric acid (37%) with 7 mL MilliQ water and dissolving 0.2 grams 1,10-Phenanthroline monohydrate (Sigma-Aldrich, 99%) in it. The complexing reagent was shaken, and 10 mL of MilliQ water was added.

An acetate buffer was prepared by mixing 3 mL MilliQ water with 12 mL acetic acid (50 mM) and dissolving 5 grams ammonium acetate (Biosolve B.V.) into it. The acetate buffer was shaken, and MilliQ water was added up to 20 mL. The reaction reagent was prepared by mixing 5 mL of the o-Phenanthroline solution with 5 mL of the acetate buffer. To determine the ferrous iron concentration of a specific sample, 0.1 mL of the sample was pipetted into a 5 mL microtube (Eppendorf). Next, 1 mL of the complexing fluoride reagent was added, the solution was shaken, and 0.4 mL of the reaction reagent was pipetted into it. Samples were agitated, then MilliQ water was added to the samples up to 2.5 mL. A volume of 200 μL of the sample was pipetted in triplicate into a 96-well plate (SARSTEDT). Samples were incubated for five minutes, then the absorbance of the 96-well plate was measured at 510 nm (Tecan infinite M200 pro microplate reader).

### Magnetic extraction apparatus

The magnetic extraction set-up consisted of three 3 mL cuvettes (Sigma Aldrich) and a neodymium magnet (60×10×3 mm, coated with Ni-Cu-Ni, MIKEDE) with a self-made Parafilm-wrapped aluminum cover. In short, the bottom half of the magnet was tightly wrapped in two layers of aluminum foil and two layers of Parafilm. The design allowed for an easy removal and insertion of the magnet. The first cuvette was filled with 1 mL of the sample, and the second and third were filled with MilliQ water. The magnet with cover was inserted into the first cuvette twice. Magnetic material was attracted to the magnet’s cover, and the covered magnet with associated magnetic material was transferred to the second cuvette for washing. The magnet with cover and magnetic material were finally placed into the third pre-weighed cuvette. To suspend the magnetic material within the MilliQ water, an additional neodymium magnet was placed underneath the cuvette. The liquid, which was not attracted by the magnet was decanted, and the cuvettes were weighted after they had dried. The original weight of each individual cuvette was subtracted from the total weight after addition of extracted material.

### Large-scale magnetic extraction

Magnetic extractions were performed by preparing a 10 g/L solution of previously sieved (<63 μm) EAC-1, JSC-2A, or JSC-Mars1 regolith simulants with TSB medium. An overnight culture of *Shewanella* was added to an O.D._600_ of 0.05 and grown for 168h at 30°C with shaking at 120 rpm. Neodymium magnets (60×10×3 mm, coated with Ni-Cu-Ni, MIKEDE) were wrapped in two layers of aluminum foil covering the ends of the magnets, after which the foil-tipped magnets were wrapped completely in a layer of Parafilm. After 168-hour incubation, the wrapped magnet was placed into the solution, the flask was swirled five times, the material was allowed to settle for 30 seconds, and the magnet was extracted again by using rubber tweezers and additional neodymium magnets. The extracted magnet was washed in a 50 mL Falcon tube containing MilliQ water. Scissors were used to cut open the middle region of the Parafilm magnet-cover, the magnet was removed from the cover, and the cover was rinsed into an empty Petri dish. This process was repeated until the amount of extracted material per magnet was so little that continuation would not result in an effectively higher yield. The non-magnetic material was collected by rinsing the remaining material out of the flask into an additional Petri dish.

### X-ray fluorescence measurements (XRF)

A pressed powder tablet was prepared by adding 0.25 g Boreox (FLUXANA) to 1 g of the tested regolith simulant (untreated and treated JSC-2A, JSC-Mars1, or EAC-1). The mixture was milled using a malachite mortar until a uniform mixture was achieved. About 5 g of Boreox was added to a hollow metal cylinder and pressed on a hydraulic press up to 10 kPa/cm^2^ (P/O/Weber Laborpresstechnik). The mixture of regolith and Boreox was added to the metal cylinder and pressed up to 250 kPa/cm^2^. The pressed tablet was analyzed with an X-ray fluorescence spectrometer (Axios Max WD-XRF, Malvern Panalytical Ltd). Analysis of the XRF data was performed with SuperQ5.0i/Omnian software.

### X-ray diffraction measurements (XRD)

Approximately 100 mg of regolith simulant was equally distributed over a silicon crystal sample holder (Si510 zero-background wafer). XRD measurements were taken using a diffractometer (D8 Advance, Bruker-AXS) with Bragg-Brentano geometry and a Lynxeye position-sensitive detector. Cu Kα radiation was used at under 45 kV and 40 mA, with a scatter screen height of 5 mm. The sample was scanned with an X-ray beam varying from 8 ° - 110 °, with a step size of 0.021 ° * 2θ (in which θ is the angle between the incident ray and the surface of the sample) and a counting time of 1 second per step. Analysis of the XRD data was performed with Bruker–AXS software DiffracSuite.EVA v 5.0.

### X-ray photoelectron spectrometry (XPS)

Samples were analyzed with X-ray photoelectron spectroscopy (XPS) on a ThermoFisher K-Alpha system using Al Kα radiation with a photon energy of 1486.7 eV. Powder samples were immobilized onto a copper tape (Plano GmbH, G3397) and were loaded into the XPS chamber without further purification. Iron high-resolution XPS spectra were acquired using a spot size of 400 μm, 50 eV pass energy, and 0.1 step size, averaging 50 scans from 705 eV to 740 eV with charge neutralizing. The peaks were calibrated for the C 1s peak at 285 eV. The background was subtracted using the “smart” function of the ThermoFisher Advantaged software.

### Lithography-based ceramic printing

The regolith simulant JSC-2A was mixed with 25.8 wt% of a photocurable organic binder (Lithoz) to prepare a printing feedstock. 3D printed pre-sintered green parts were produced on a CeraFab 7500 using vat photopolymerization. A light engine (based on LEDs) with a digital micromirror device was used to selectively harden the previously prepared regolith simulant containing feedstock layer by layer into desired shape based on a given STL file. The 3D printed green parts were subsequently dried and sintered for 2 hours at 1050°C and 1100°C for the untreated and treated samples, respectively.

### Ultimate Compressive Strength of 3D prints

Compressive tests were performed to determine the ultimate compressive strength of the 3D printed materials (Instron ElectroPuls E10000 Linear-Torsion) under environmental conditions of 20°C and a relative humidity of approximately 50%. DIN 51104 standards were followed with two exceptions: no intermediate plate was used between the instrument and the samples, and the specimens were not fully compliant to the standard since the ends of the 3D prints were not perfectly flat. A constant deformation rate of 0.5 mm/min was used. The ultimate compressive strength was calculated using the following formula: σ = F* / A_0_, where σ is the ultimate compressive strength, F^*^ is the force applied to the sample just before it cracks, and A0 is the initial area of the sample.

## Acknowledgments

Our thanks to Florian Ertl for the 3D printing work performed and to Stan Brouns and Hubertus J. E. Beaumont for providing access to their laboratories. This work was supported by The Netherlands Organization for Scientific Research (NWO/OCW), as part of the Frontiers of Nanoscience program.

## Notes

### Competing Interest Statement

The authors have declared no competing interest.

